## Supplementary Material for "Heritability of Urinary Amines, Organic Acids, and Steroid Hormones in Children"

### **Supplementary Results**

#### Sensitivity and sex- and age-stratified analyses for urinary metabolites

First, we repeated the mixed-effect analyses after dichotomizing the age variable (Age<10 vs. Age≥10). Dichotomizing the age variable had little influence on the association of sex with metabolites (*r_beta_* = 0.99, *p* < 2.902x10^-161^, **Figure S3a**), all metabolites associated with sex in the model with continuous age also show a significant association after dichotomization, though we observe an additional association with sex for 3-hydroxybutyric acid (**Figure S2**, **Table S1**, **Table S9**). Similarly, the regression estimates for continuous versus dichotomized age also had near perfect correlations (*r_beta_* = 0.99, *p* < 1.01x10^-80^, **Figure S3b**), though the associations for 3-hydroxyisovaleric acid, beta-alanine, glycine, hydroxylysine, and L-lysine as observed with continuous age were no longer significant when dichotomizing age, and the association with glycolic acid was only observed after dichotomizing age (**Figure S2**, **Table S1**, **Table S9**).

Sex-stratified analyses showed that 65 metabolites with a significant association with continuous age in males also had significant associations in females (**Figure S2**, **Table S10**). For 5 amines, one organic acid, and 2 LC-MS steroids, we observed significant associations in males but not in females, and for one amine and one organic acid we observed significant association in females but not in males. Thus sex-stratified analyses showed that age mostly influenced the same metabolites in males and females, with on average less pronounced age effects observed in females, though the association strength in males and females was highly correlated (*r_beta_* = 0.96, *p* <1.07x10^-47^, **Figure S3c**). We also observed strong correlations among the strength of the associations for continuous age as observed in males and females separately with the strength of the associations for continuous age in males and females combined (*r* = 0.99, *p* <6.42x10^-76^, **Figure S3d**, *r* = 0.99, *p* <6.36x10^-69^, **Figure S3f**, respectively), and with the strength of the associations for dichotomized age in males and females combined (*r* = 0.99, *p* <2.11x10^-68^, **Figure S3e**, *r* = 0.98, *p* <5.88x10^-59^, **Figure S3g**, respectively). Repeating the sex-stratified analyses after dichotomization of age, showed 47 metabolites with significant age associations in males and females, and showed high correlations in the strength of association between males and females and between dichotomized and continuous results (**Figure S2**, **Table S11**, **Figure S3h-n**).

Stratifying analyses by age (Age<10 vs. Age≥10) reduced the number of metabolites with significant associations with sex (**Figure S2**, **Table S12**). For 10 metabolites we observed significant associations with sex in both age groups, 3-methoxytyrosine only showed a significant association with sex in the younger age group, while L-phenylalanine, alpha-cortolone, and dehydroepiandrosterone sulfate (DHEA-S) only showed significant sex associations in the older age group. Overall, the effect size estimates were highly similar in both age groups (*r* = 0.90, *p* <2.37x10^-32^, **Figure S3o**), and comparing the effect sizes per age group to those observed for the continuous and dichotomized estimates in the combined age groups showed high concordance (**Figure S3p-s**).

We replicated the significant sex and age associations for the amines, organic acids, and LC-MS steroids in 179 children of the LUMC-Curium cohort (*M* age = 10.2, *SD* age = 1.8, age range: 6.3-13.4, 25.1% females). The associations of continuous age with 30 metabolites, dichotomized age with 4 metabolites, continuous age in females with 3 metabolites, and of sex in the older (Age≥10) age group with one metabolite remained significant in the replication cohort (**Figure S2**, **Table S2** and **Tables S13-S16**). While the number of replicated associations for sex and age was low, we observed high correlations among the strength and direction of association between the discovery and replication results (**Figure S1**).

### **Supplementary Figures**

#### **Figure S1.** Correlations of the strength of association (beta coefficients) for sex and age with metabolites between the NTR-ACTION discovery cohort and LUMC-Curium ACTION replication cohort.

**(a)** Correlation of the beta coefficients of the sex associations in the discovery vs replication cohorts in the model with continuous age. **(b)** Correlation of the beta coefficients of the continuous age associations in the discovery vs replication cohorts. **(c)** Correlation of the beta coefficients of the sex associations in the discovery vs replication cohorts in the model with dichotomized age. **(d)** Correlation of the beta coefficients of the dichotomized age associations in the discovery vs replication cohorts. **(e)** Correlation of the beta coefficients of the continuous age associations in the sex-stratified models for males in the discovery vs replication cohorts. **(f)** Correlation of the beta coefficients of the continuous age associations in the sex-stratified models for females in the discovery vs replication cohorts. **(g)** Correlation of the beta coefficients of the dichotomized age associations in the sex-stratified models for males in the discovery vs replication cohorts. **(h)** Correlation of the beta coefficients of the dichotomized age associations in the sex-stratified models for females in the discovery vs replication cohorts. **(i)** Correlation of the beta coefficients of the sex associations in the age-stratified models for younger children (Age<10) in the discovery vs replication cohorts. **(j)** Correlation of the beta coefficients of the sex associations in the age-stratified models for older children (Age≥10) in the discovery vs replication cohorts. Significant associations of sex or age with metabolites in both the discovery and replication cohorts are depicted in black, and significant associations of sex or age with metabolites only in the discovery depicted in grey. All association results for all models in the discovery are reported in **Table S1** and **Tables S9-S12** and all association results for all models in the replication are reported in **Table S2** and **Tables S13-S16**.

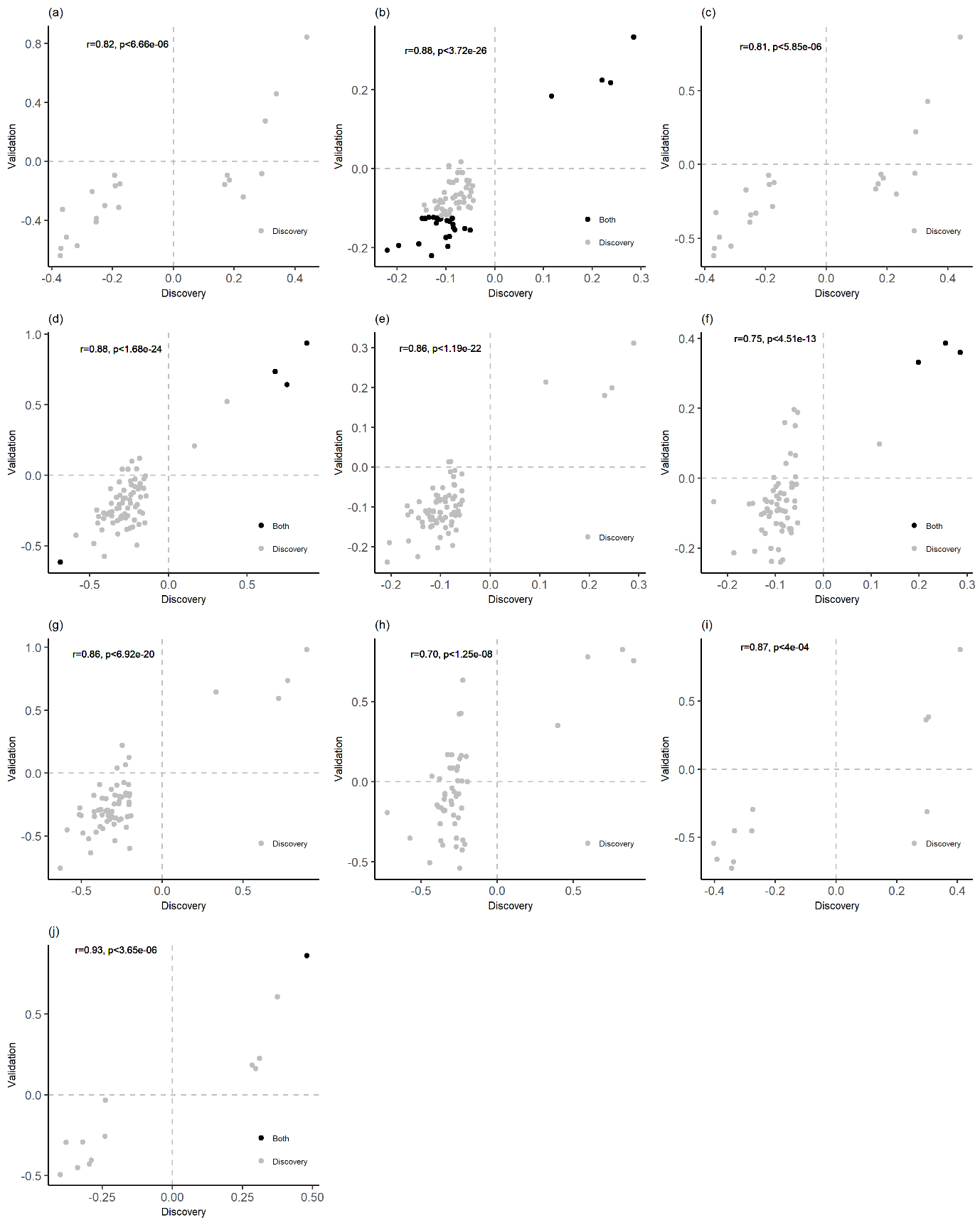

#### **Figure S2.** Comparison of the associations of sex and age with metabolites in the main, sensitivity, and stratified analyses in the NTR and LUMC-Curium ACTION cohorts.

The vertical axis lists the metabolites and the horizontal axis depicts the associations for sex or age in the various models as performed in the discovery NTR-ACTION cohort (NTR) or in the replication LUM-Curium ACTION cohort (Curium). The ‘C’ suffix indicates the model included age as a continuous variable, while the D suffix indicates the model included age as a dichotomized variable (Age<10 vs. Age≥10). The ‘boys’ and ‘girls’ suffixes indicate these are the associations as obtained from the sex-stratified models, and the ‘<10y’ and ‘>-10y’ suffixes indicate that these are the associations as obtained from the age-stratified analyses. All associations marked with a star (*) where significant after correcting for multiple testing (see **Methods**). The color gradient indicates the strength of the associations (beta coefficient [B]), where dark blue indicates strong negative associations and dark red indicates strong positive associations. All association results for all models in the discovery are reported in **Table S1** and **Tables S9-S12** and all association results for all models in the replication are reported in **Table S2** and **Tables S13-S16**.

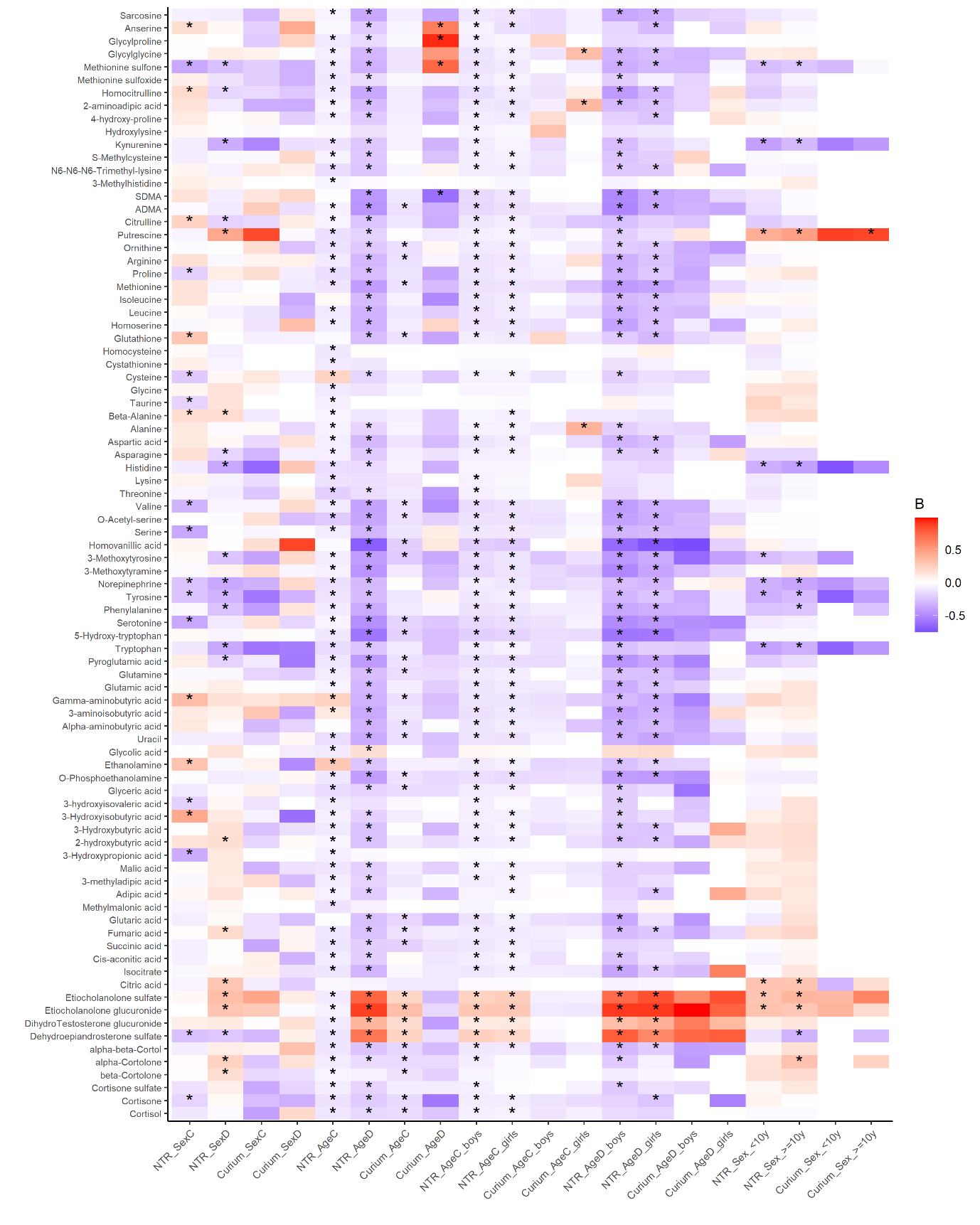

#### **Figure S3.** Correlations of the strength of association (beta coefficients) between the different association models for sex and age with metabolite levels in the NTR-ACTION discovery cohort.

**(a)** Correlation of the beta coefficients of the sex associations in the model with continuous age vs dichotomized age. **(b)** Correlation of the beta coefficients of the age associations in the model with continuous age vs dichotomized age. **(c)** Correlation of the beta coefficients of the continuous age associations in the sex-stratified models males vs females. **(d)** Correlation of the beta coefficients of the continuous age associations in the sex-stratified model for males vs the continuous age associations in males and females. **(e)** Correlation of the beta coefficients of the continuous age associations in the sex-stratified model for males vs the dichotomized age associations in males and females. **(f)** Correlation of the beta coefficients of the continuous age associations in the sex-stratified model for females vs the continuous age associations in males and females. **(g)** Correlation of the beta coefficients of the continuous age associations in the sex-stratified model for females vs the dichotomized age associations in males and females. **(h)** Correlation of the beta coefficients of the dichotomized age associations in the sex-stratified model for males vs females. **(i)** Correlation of the beta coefficients of the continuous vs dichotomized age associations in the sex-stratified model for males. **(j)** Correlation of the beta coefficients of the continuous vs dichotomized age associations in the sex-stratified model for females. **(k)** Correlation of the beta coefficients of the dichotomized age associations in the sex-stratified model for males vs the continuous age associations in males and females. **(l)** Correlation of the beta coefficients of the dichotomized age associations in the sex-stratified model for males vs the dichotomized age associations in males and females. **(m)** Correlation of the beta coefficients of the dichotomized age associations in the sex-stratified model for females vs the continuous age associations in males and females. **(n)** Correlation of the beta coefficients of the dichotomized age associations in the sex-stratified model for females vs the dichotomized age associations in males and females. **(o)** Correlation of the beta coefficients of the sex associations in the age-stratified models Age<10 vs Age≥10. **(p)** Correlation of the beta coefficients of the sex associations in the age-stratified models Age<10 vs the sex associations in the models including continuous age associations in Age<10 and Age≥10. **(q)** Correlation of the beta coefficients of the sex associations in the age-stratified models Age<10 vs the sex associations in the models including dichotomized age associations in Age<10 and Age≥10. **(r)** Correlation of the beta coefficients of the sex associations in the age-stratified models Age≥10 vs the sex associations in the models including continuous age associations in Age<10 and Age≥10. **(s)** Correlation of the beta coefficients of the sex associations in the age-stratified models Age≥10 vs the sex associations in the models including dichotomized age associations in Age<10 and Age≥10. Colors indicate whether no significant associations were found in either model (grey), in both models (black), only before dichotomization of the age variable (green), only after dichotomization of the age variable (yellow), only in males (dark blue), only in females (red), only in Age<10 (orange), or only in Age≥10 (light blue). All association results for all models in the discovery are reported in **Table S1** and **Tables S9-S12**.

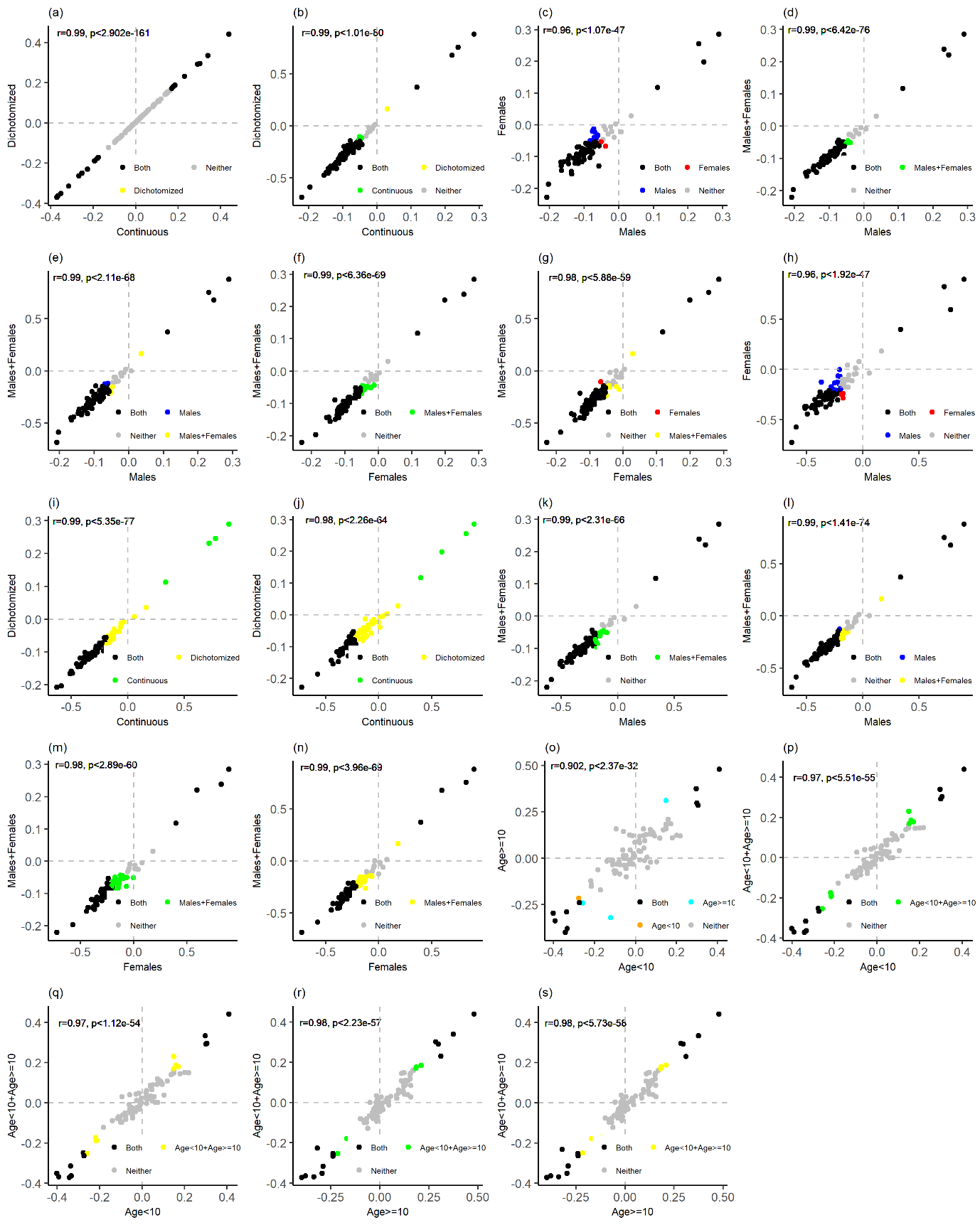

### **Supplementary Tables**

#### **Table S7.** Participant characteristics for the NTR-ACTION cohort by aggression concordance at the time of invitation into the ACTION Biomarker Study.

|  | **Concordant low-low aggression pairs** | | | | **Discordant low-high aggression pairs** | | | | | **Concordant high-high aggression pairs** | | | |
| --- | --- | --- | --- | --- | --- | --- | --- | --- | --- | --- | --- | --- | --- |
|  | **N (N pairs)** | **Mean (SD) [range] age** | **N (%) females** | **Mean (SD) [range] AGG^1^** | **N (N pairs)** | **Mean (SD) [range] age** | **N (%) females** | **Mean (SD) [range] AGG^1^** | | **N (N pairs)** | **Mean (SD) [range] age** | **N (%) females** | **Mean (SD) [range] AGG^1^** |
|  |  |  |  |  |  |  |  | **Low^2^** | **High^2^** |  |  |  |  |
| Total | 585 (290) | 9.4 (1.8) [5.7 - 12.6] | 310 (53%) | 2.8 (3.8) [0 - 25] | 362 (180) | 10.1 (1.7) [6.1 - 12.7] | 161 (44.5%) | 4.4 (4.5) [0 - 19] | 6.3 (5.8) [0 - 30] | 353 (175) | 9.5 (1.8) [6 - 12.9] | 155 (43.9%) | 7.5 (6) [0 - 30] |
| MZ | 454 (226) | 9.1 (1.9) [6.1 - 12.6] | 240 (52.9%) | 2.6 (3.7) [0 - 21] | 295 (147) | 10.2 (1.6) [6.1 - 12.7] | 124 (42%) | 4.5 (4.7) [0 - 20] | 4.9 (5.1) [0 - 22] | 319 (158) | 9.6 (1.8) [6 - 12.9] | 142 (44.5%) | 7.3 (5.9) [0 - 30] |
| DZ | 131 (64) | 10.3 (1.3) [5.7 - 12] | 70 (53.4%) | 3.4 (4.2) [0 - 25] | 67 (33) | 9.6 (1.8) [6.2 - 11.9] | 37 (55.2%) | 7.5 (5.2) [0 - 21] | 8.4 (6.1) [0 - 30] | 34 (17) | 9.2 (1.8) [6.2 - 11.4] | 13 (38.2%) | 9.2 (6.3) [0 - 24] |

Notes: AGG, aggression; NTR, Netherlands Twin Register, MZ, monozygotic twin pairs; DZ, dizygotic twin pairs.

^1^ Measured with the mother-rated Aggressive Behavior syndrome scale of the Achenbach System of Empirically Based Assessment (ASEBA) Child Behavior Checklist (CBCL) at time of biological sample collection.

^2^ In the twin pairs discordant for aggressive behavior at the time of invitation into the ACTION Biomarker Study (Hagenbeek et al. 2020), we provide the mean, standard deviation (SD), and range of the mother-rated ASEBA CBCL Aggressive Behavior syndrome scale as measured at the time of biological sample collection separately for the twin with a low and a high aggression score at time of invitation into the ACTION Biomarker Study.

**Table S8.** Internal standards used in the LC-MS amine platform.

| Asn_C13N15 | Asp_C13N15 | L-ornithine-3,3,4,4,5,5,-d6 |
| --- | --- | --- |
| L-NT-methyl-d3-L-histidine | Glu_C13N15 | Lys C13N15 |
| Ser_C13N15 | Beta-alanine-2,2,3,3,-d4 | Tyr_C13N15 |
| Gln_C13N15 | Thr_C13N15 | L-Methionine_C13N15 |
| Arg_C13N15 | Ala_C13N15 | Val_C13N15 |
| Gly_C13N15 | Phe_C13N15 | Trp_C13N15 |
| Histamine-α,α,β,β-d4 2HCl | L-2-aminobutyric acid-d6 acid | 2-(4-hydroxy-3-methoxyphenyl) ethyl-1,1,2,2-d4-amine |
| L-Ile_C13N15 | Leu_C13N15 |  |
